# Soil and vegetation conditions changes following the different sand dune restoration measures on the Zoige Plateau

**DOI:** 10.1101/627091

**Authors:** Jiufu Luo, Dongzhou Deng, Li Zhang, Xinwei Zhu, Dechao Chen, Jinxing Zhou

## Abstract

Restoration of alpine sand dunes has been increasingly attracting the attention of ecologists due to their difficulty and importance among the mountain-river-forest-farmland-lake-grass system (referred as meta-ecosystem) restoration. Alpine sand dunes are suffered from unstable soil and lack of plants. Efficient restoration measures are vital to guide the sand dune restoration. Whether the engineering materials co-applied with seeding could achieve considerable restoration in such areas? Here, sandbag and wicker as environmental friendly materials combined with *Elymus nutans* seeding were implemented on the Zoige Plateau sand dune, comparing with the ‘control’ treatment that only seeding. We assessed the topsoil conditions by sampled the surface soil and measured the water capacity and nutrients. We also utilized interspecific relationship and population niche to analyse the plant community structure variances among different restoration measures. Results showed that the soil conditions got clearly improved in sandbag area than that in wicker area when compared with that in control area. The community in control area was the least structured, while the species showed the closest related in sandbag area. In addition, average population niche overlap showed a control (0.26) < wicker (0.32) < sandbag (0.39) ranking. Thus, we suggested that sandbag or wicker co-applied with indigenous grass seeding is a practical and quick restoration approach in alpine sand dunes, and the sandbag may surpasses the wicker. Moreover, soil amending measures including nutrient improvement, and microbial fertilizer addition may further accelerate sand dune restoration.

## 1. Introduction

The terrestrial ecosystem has experienced increasingly severe land degradation and desertification [1]. Desertification threatens the ecological safety and its restoration is one of the vital elements in the mountain-river-forest-farmland-lake-grass system (referred as meta-ecosystem) restoration [2, 3]. The long-termed complex causes (e.g., overgrazing, climate change) accelerate desertification process and the sand dunes are expanding continuously on the Zoige Plateau [4-6]. Sand dunes are generally covered by nutrient devoid sandy soil that lacks favourable properties for plant growth and is prone to wind erosion [7]. The expanded sand dunes destroy fertile land and welfare of local populations. For example, they threaten livestock productivity, society development, ecological civilization, household income or human beings health [8-10]. Hence, sand dunes restoration is crucial in ensuring the conservation and sustainable development of the Zoige Plateau and meeting aspiration for better living standards in local people.

Decades worth of works have been conducted on sand dunes restoration [11, 12]. Mechanical sand barriers as the classical and simplest measure have been shown to contribute to reduce sand dunes mobility [13]. Stone or straw checkerboard barriers have been utilized to fix active sand dunes along the railway in alpine sandy land, such as Baotou-Lanzhou Railway and the eastern shore of the Qinghai Lake [14]. However, the single use of types of barriers is non-optimal choice in alpine sand dune restoration. They were often buried by sand sediments due to lack of vegetation cover eventually, in addition, traditional mechanical materials were much expensive and hard to operate [13].

Vegetation restoration is an important objective during sand dune restoration [15, 16]. It has been proved to be practical in decreasing wind velocity and increasing soil nutrients in sand dune to accelerate vegetation restoration [17-19]. Given the poor seed bank in active sand dunes, natural vegetation restoration is almost not feasible [20, 21], so that of indigenous seeds application is an inevitable method to improve seed bank [22, 23]. For example, marram grass (*Ammophila arenaria*) was one of an optimal species for sand dune restoration in Ille et Vilaine, France [11]. ‘Tree-screens’ and ‘shelter-belt’ plantations of the Thar Desert in India were launched and achieved great ecological benefits in vegetation cover [17]. Sahara mustard (*Brassica tournefortii*) usually occupied dominated status in the drier sand dune region, and the flourishing weed also could control sand dune in the semi-arid regions of Inner Mongolia [24, 25]. In addition, Farmland constructed in the Mu Us Sandy Land has changed the barren desert to fertile farmland over ten-year restoration, providing compelling evidence of biotic approaches advantages in sand dune restoration [26]. Also, shrub-planting is an effective restoration measure to fix the sand dune in the semiarid Mu Us desert [27].

Although there were plenty of successful project cases of sand dunes restoration around the world previously, few of them were suitable for sand dune restoration on the alpine area. Here, we focus on two environmental friendly barrier materials (i.e., Poly Lactic Acid sandbag and *Salix paraplesia* wicker) that are easily reproducible and durable in harsh conditions. Poly Lactic Acid is hydrophilic, ultraviolet radiation resistance, and easy transportation [28], making them as optimal barrier materials in sand dune restoration on the Zoige Plateau. Meanwhile, *S. paraplesia* is widely cultivated in alpine area which makes it convenient to acquire wicker materials. These two materials combined with indigenous grass (*Elymus nutans*) were used with expectation to fix the active sand dune on the alpine sand dunes. Our objectives were compare the different restoration approaches’ effects on the alpine sand dunes and expect to provide a suitable strategy for alpine sand dune restoration.

## 2. Materials and Methods

### 2.1. Study area

The study area is located in Xiaman, Assi Township, on the Zoige Plateau (3,486 m asl.), which is characterized by an alpine continental monsoon climate with a pronounced winter season. The annual average temperature is 2.5°C and the average annual rainfall is 520 mm. The maximum wind speed is up to 36 m/s, with northwest prevailing winds [9]. The soil is dominated by alpine or subalpine meadow soil, with marshlands distributed throughout. Over the years, the land degradation process has increased greatly. Thus it led to types of degraded landscapes, form a large area of active sand dune. The active sand dune has caused serious threatens to ecological safety.

### 2.2. Field investigation design and sampling

We established a restoration demonstration zone in a 10 ha degraded land in which active sand land occupied more than 55%. This area is a typical degraded alpine land on the Zoige Plateau. We employ sandbag and wicker as sand barrier materials combined with *Elymus nutans* (60 kg hm^-2^) sowing in the study area (referred as “sandbag” and “wicker”). The sand dune that only implemented sowing was set as ‘control’. Thus, three restoration measures areas consist of control, wicker, and sandbag area were organized in a randomized block design in the restoration demonstration zone. The barrier checkerboard was 2.0 m × 2.0 m × 0.3 m. (Fig 1)

**Fig 1.**
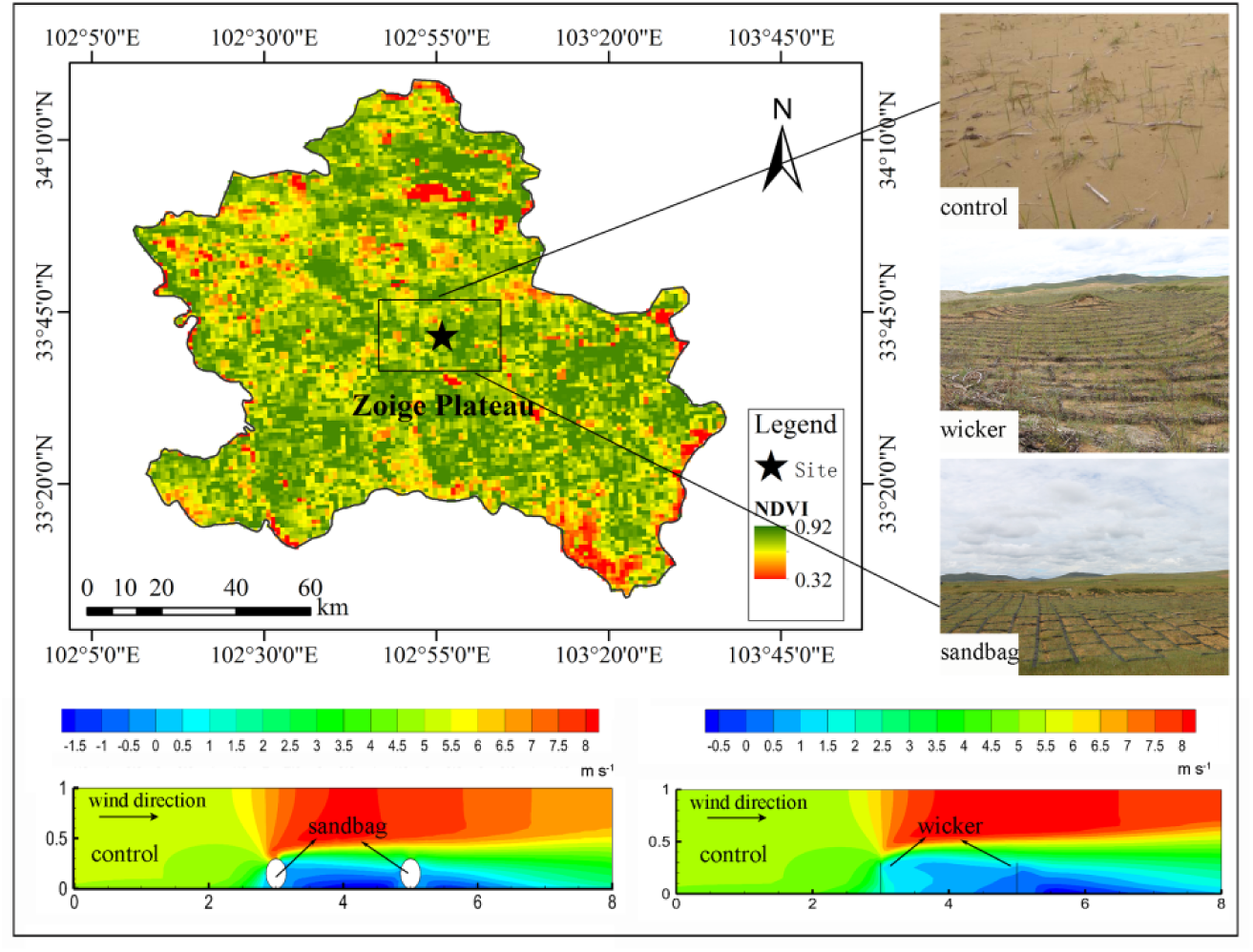
Demonstration area and speed flow distribution characteristics of different restoration measure on the Zoige Plateau.

108 quadrats were randomly investigated in three restoration areas at the third growth season after the restoration measures were implemented. Parameters of plant taxa, natural plant height, species cover and stem number were recorded. Surface soil in each quadrat was sampled by soil core, sieved (< 2 mm) to filter out gravel or plant roots and divided into three subsamples. One was saved in a refrigerator (4°C) for microbial biomass carbon (MBC) and microbial biomass nitrogen (MBN) determination by the chloroform fumigation-incubation method co-applied with an N/C Analyser (multi N/C® 3100 TOC, analytikjena, Germany); one was air-dried to measure total soil organic carbon (thereafter SOC) by titrimetry, total soil nitrogen (TN), total soil phosphorus (TP), soil ammonium nitrogen (AN), soil nitric nitrogen (NN), soil available phosphorus (AP) by Smartchem Discrete Auto Analyzer (Smartchem 200, AMS/Westco, Italy); and one was used for soil moisture determination using the gravimetrical method by drying at 105°C.

### 2.3. Data analysis

The species importance value (*IV*) was calculated using the following equation:

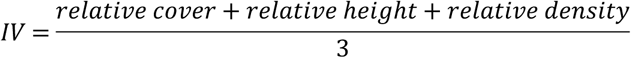

The Jaccard interspecific association (*JI*) test was conducted based on the 2 × 2 contingency tables by the plant investigation data (Table 1).

**Table 1.** Illustration of the 2×2 contingency tables.

|  | Quadrats number of species j appeared | Quadrats number of species j absented |
| --- | --- | --- |
| Quadrats number of species i appeared | a | b |
| Quadrats number of species i absented | c | - |

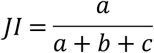

The *JI* value was classified into 4 grades: none association, 0 ≦ *JI* ≦ 0.25; weak association, 0.25 < *JI* ≦ 0.5; middle association, 0.5 < *JI* ≦ 0.75; and song association, 0.75 < *JI* ≦ 1.0 [29].

Furthermore, the Spearman rank correlation (*r*(*i, k*)) was tested to assess the interspecific correlation degree.

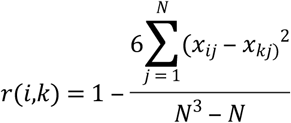

Here, *r(i, k)* is the correlation coefficient between species *i* and *k, x*_*ij*_ and *x*_*kj*_ are the importance values of species *i* and *k* in quadrat *j*.

Additionally, niche theory has been widely used in the study of plant community ecology [30]. Niche breadth and overlap are important indices to further quantify the resource utilization efficiency and competition/coexistence of different populations [31-33]. Shannon-Wiener niche breadth (*B*_*i*_) was calculated following Colwell & Futuyma [34] and The Pianka niche overlap (*O*_*ik*_) was calculated using the following equation [35]:

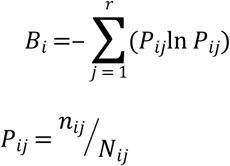

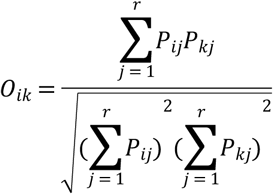

Here, *P*_*ij*_ and *P*_*kj*_ are a proportion of quadrat *j* among the total quadrats occupied by species *i* and *k*; *r* is the total number of quadrats. The n_*ij*_ is the importance values of species *i* in quadrat *j* and *N*_*ij*_ = Σ*n*_*ij*_.

The soil metric and species richness data were calculated using MS Excel 2010, and statistical analyses were performed using SPSS Statistics 20.0 (*P*<0.05) (SPSS Inc., Chicago, IL, US). The soil condition graphs were run with OriginPro 2016 (OriginLab Corporation, Northampton, MA, US). The map was created with ArcGIS v10.2. The speed flow distribution characteristics were simulated by Gambit 2.4, Fluent 16.0, and Tecplot 360. The niche overlap matrix diagram was run with the ‘Lattice’ package in R. The Jaccard interspecific association graphs, niche overlap matrix diagrams, and field experimental site pictures were merged by Adobe Photoshop CS6 v6.0.335.0.

## 3. Results

### 3.1. Plant composition and soil conditions in different restoration area

We recorded 9, 12, and 10 plant species in the sandbag, wicker and control area, respectively. The vegetation cover increased by 161.99% in wicker area and 331.67% in sandbag area compared with that in control (*P*<0.05). The same species importance values varied among the different restoration area. The *E. nutans* occupied the dominant position in all restoration area, especially in the sandbag area in which its importance value was up to 52.89. (Table 2)

**Table 2.** Plant composition, importance value (*IV*), niche breadth (*Bi*), and vegetation cover (VC) in different restoration area.

| Sandbag |  |  |  | Wicker |  |  |  | Control |  |  |  |
| --- | --- | --- | --- | --- | --- | --- | --- | --- | --- | --- | --- |
| Species | *IV* | *Bi* | *VC* | Species | *IV* | *Bi* | *VC* | Species | *IV* | *Bi* | *VC* |
| *En.* | 52.89 | 1.55 |  | *En.* | 32.03 | 1.54 |  | *En.* | 39.57 | 1.55 |  |
| *Cm.* | 17.04 | 1.54 |  | *Ls.* | 19.43 | 1.48 |  | *Cm.* | 19.64 | 1.42 |  |
| *Ls.* | 10.86 | 1.50 |  | *Hb.* | 12.21 | 1.50 |  | *Kr.* | 17.77 | 1.46 |  |
| *Hb.* | 8.19 | 1.41 |  | *Of.* | 10.98 | 1.40 |  | *Ls.* | 7.48 | 1.17 |  |
| *Of.* | 7.78 | 1.39 | 19.08 | *HL.* | 7.61 | 1.40 | 11.58 | *Pb.* | 4.38 | 0.91 | 4.42 |
| *Fo.* | 1.43 | 0.69 | ±0.77c | *Cm.* | 7.48 | 1.11 | ±0.70b | *Hb.* | 4.20 | 1.07 | ±0.16a |
| *Dh.* | 1.27 | 0.82 |  | *Kr.* | 5.52 | 1.24 |  | *HL.* | 2.28 | 0.76 |  |
| *Ms.* | 0.31 | 0.48 |  | *Am.* | 1.27 | 0.58 |  | *Of.* | 2.03 | 0.48 |  |
| *Od.* | 0.23 | 0.30 |  | *Ps.* | 1.24 | 0.38 |  | *Dh.* | 1.82 | 0.69 |  |
|  |  |  |  | *Ms.* | 0.87 | 0.77 |  | *Ms.* | 0.82 | 0.48 |  |
|  |  |  |  | *Sc.* | 0.84 | 0.48 |  |  |  |  |  |
|  |  |  |  | *Ap.* | 0.54 | 0.48 |  |  |  |  |  |

Note: *En.*, *Elymus nutans*; *Cm.*, *Carex moorcroftii*; *Kr.*, *Kobresia robusta*; *Ls.*, *Ligusticum scapiforme*; *Pb.*, *Potentilla bifurca*; *Hb.*, *Heteropappus bowerii*; *HL.*, *Hypochaeris leptocarpum*; *Of.*, *Oxytropis falcate*; *Dh.*, *Dracocephalum heterophyllum*; *Ms.*, *Microula sikkimensis*; *Am.*, *Artemisia macrocephala*; *Ps.*, *Polygonum sibiricum*; *Sc.*, *Salsola collina*; *Ap.*, *Axyris prostrata*; *Fo.*, *Festuca ovina*; *Od.*, *Oxytropis densa*.

The soil water capacity and nutrient metrics increased greatly in the area where the sand barriers were implemented (*P*<0.05). The atomic ratios of SOC: TN, SOC: TP, TN: TP, and MBC: MBN varied in different restoration area. Comparing with the control, MBC: MBN ratios decreased a lot in sandbag area and wicker area, while the SOC: TP and TN: TP ratios increased. In more detail, the SOC: TN and MBC: MBN were only 11.67± 1.46 and 10.57± 0.21 in sandbag area, which was lower than that in wicker and control areas; the MBC: MBN in sandbag area was less than one-half of that in control area. The TN: TP and the SOC: TP ratios also were the highest in sandbag area while lowest in control area. (Table 3; Fig 2)

**Table 3.** The soil conditions variances in different restoration area (*P* < 0.05).

|  | Moisture/ % | TN /<br>g kg<sup>-1</sup> | TP/<br>g kg<sup>-1</sup> | AN/<br>mg kg<sup>-1</sup> | NN/<br>mg kg<sup>-1</sup> | AP/<br>mg kg<sup>-1</sup> | SOC/<br>g kg<sup>-1</sup> | MBC/<br>mg kg<sup>-1</sup> | MBN/<br>mg kg<sup>-1</sup> |
| --- | --- | --- | --- | --- | --- | --- | --- | --- | --- |
| **Sandbag** | 4.11 a<br>±0.02 | 0.19 a<br>±0.02 | 0.30 a<br>±0.00 | 24.93 a<br>±2.04 | 7.68 a<br>±0.24 | 84.05 a<br>±0.24 | 2.19 a<br>±0.10 | 65.83 a<br>±0.31 | 6.23 a<br>±0.14 |
| **Wicker** | 3.60 b<br>±0.02 | 0.10 b<br>±0.01 | 0.33 a<br>±0.01 | 22.19 a<br>±1.24 | 5.78 b<br>±0.50 | 73.37 b<br>±0.34 | 1.66 b<br>±0.09 | 52.37 b<br>±0.17 | 3.53 b<br>±0.14 |
| **Control** | 2.81 c<br>±0.08 | 0.06 b<br>±0.00 | 0.22 b<br>±0.02 | 13.39 b<br>±0.55 | 3.07 c<br>±0.74 | 43.80 c<br>±0.71 | 0.95 c<br>±0.03 | 15.27 c<br>±0.60 | 0.67 c<br>±0.07 |

Note: The different lowercase letters means significant difference in a certain soil condition ( $P < 0.05$ ).

**Fig 2.**
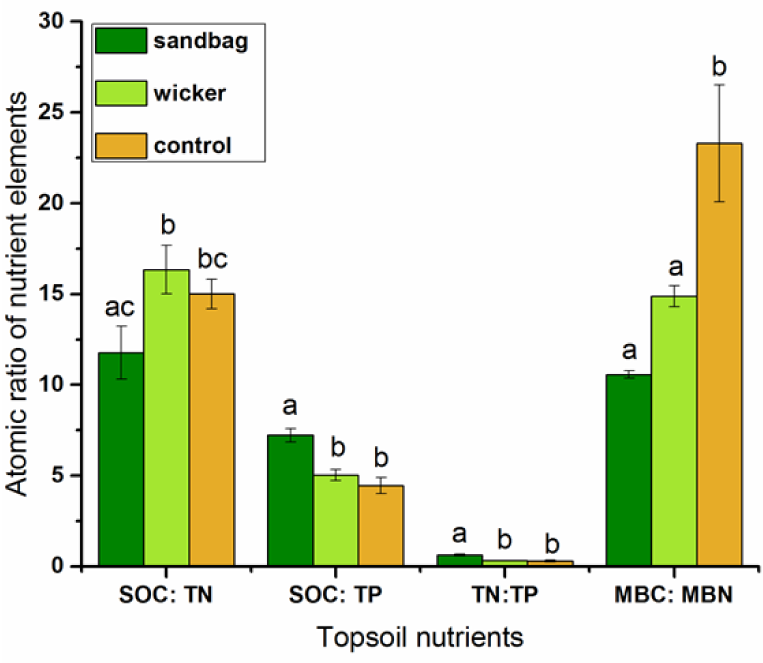
The soil nutrients atomic ratios in different restoration area. The different lowercase letters means significant difference in a certain pair-wise (*P* < 0.05).

### 3.2. Interspecific relationship in different restoration area

The summed ratio of none and weak associations were 93.33%, 83.33%, and 75.00% in control, wicker, and sandbag areas, respectively. Both of strong and middle association ratios rank was control < wicker < sandbag. The plant community was the simplest structured in the control area, and less structured in wicker area compared with that in sandbag area where the species association degree was stronger. It indicated that the community stability and community development status got better in sand barriers areas (Fig 3).

**Fig 3.**
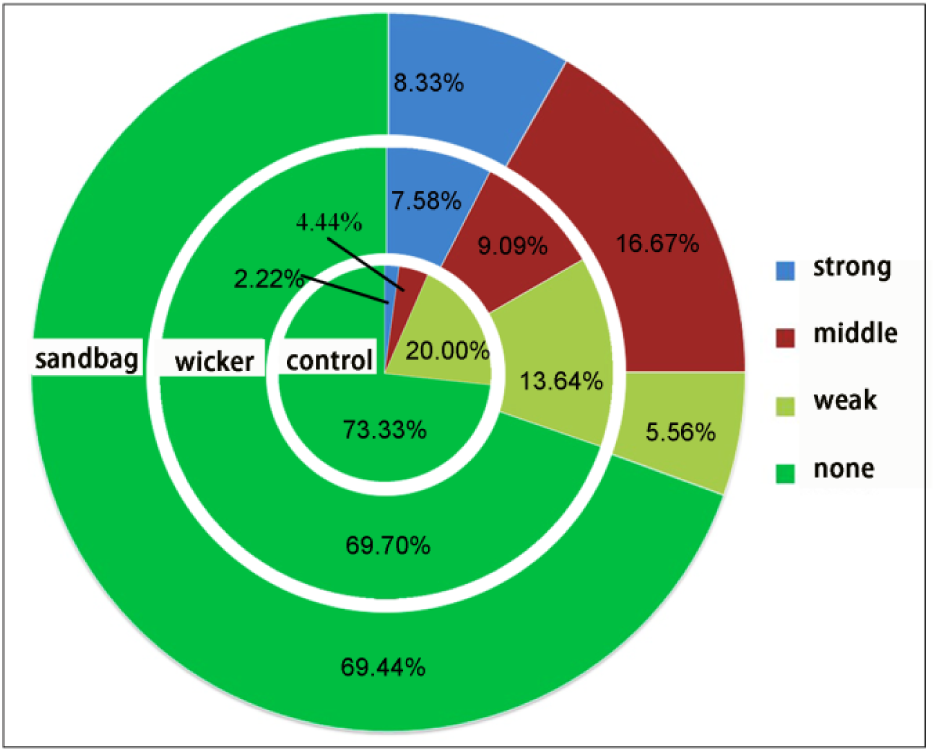
The Jaccard interspecific association (*JI*) ratios in different restoration area.

The species Spearman rank correlation indices were mostly negative in all restoration areas. *E. nutans* was negatively correlated with most of plants. The positive correlation ratio was the lowest in sandbag area. In addition, the same species pair interspecific correlation changed when the sand barriers were implemented to fix the active sand land. For example, the interspecific correlation of *E. nutans*-*C. moorcroftii* changed from greatly negative correlation to positive correlation (−0.47 (*P*<0.01) in control, −0.18 in wicker, and 0.10 in sandbag area); and the negative correlation degree of *E. nutans*-*H. bowerii* was enhanced in wicker and sandbag area (−0.08 in control, −0.34 (*P*<0.05) in wicker, and −0.47 (*P*<0.01) in sandbag area) (Tables 4-6).

**Table 4.** Spearman rank correlation between species-pair in sandbag area.

|  | *En* | *Cm* | *Ls* | *Hb* | *Of* | *Fo* | *Dh* | *Ms* |
| --- | --- | --- | --- | --- | --- | --- | --- | --- |
| *Cm* | 0.096 |  |  |  |  |  |  |  |
| *Ls* | -0.217 | -0.149 |  |  |  |  |  |  |
| *Hb* | -0.466** | -0.050 | -0.266 |  |  |  |  |  |
| *Of* | -0.006 | -0.489** | -0.318 | -0.189 |  |  |  |  |
| *Fo* | -0.055 | 0.113 | -0.528** | 0.148 | -0.081 |  |  |  |
| *Dh* | -0.339* | -0.363* | 0.629** | -0.071 | -0.246 | -0.195 |  |  |
| *Ms* | -0.470** | -0.254 | -0.442** | 0.393* | 0.395* | 0.508** | -0.147 |  |
| *Od* | -0.108 | 0.203 | -0.265 | 0.190 | -0.084 | -0.097 | -0.118 | -0.073 |

**Table 5.**
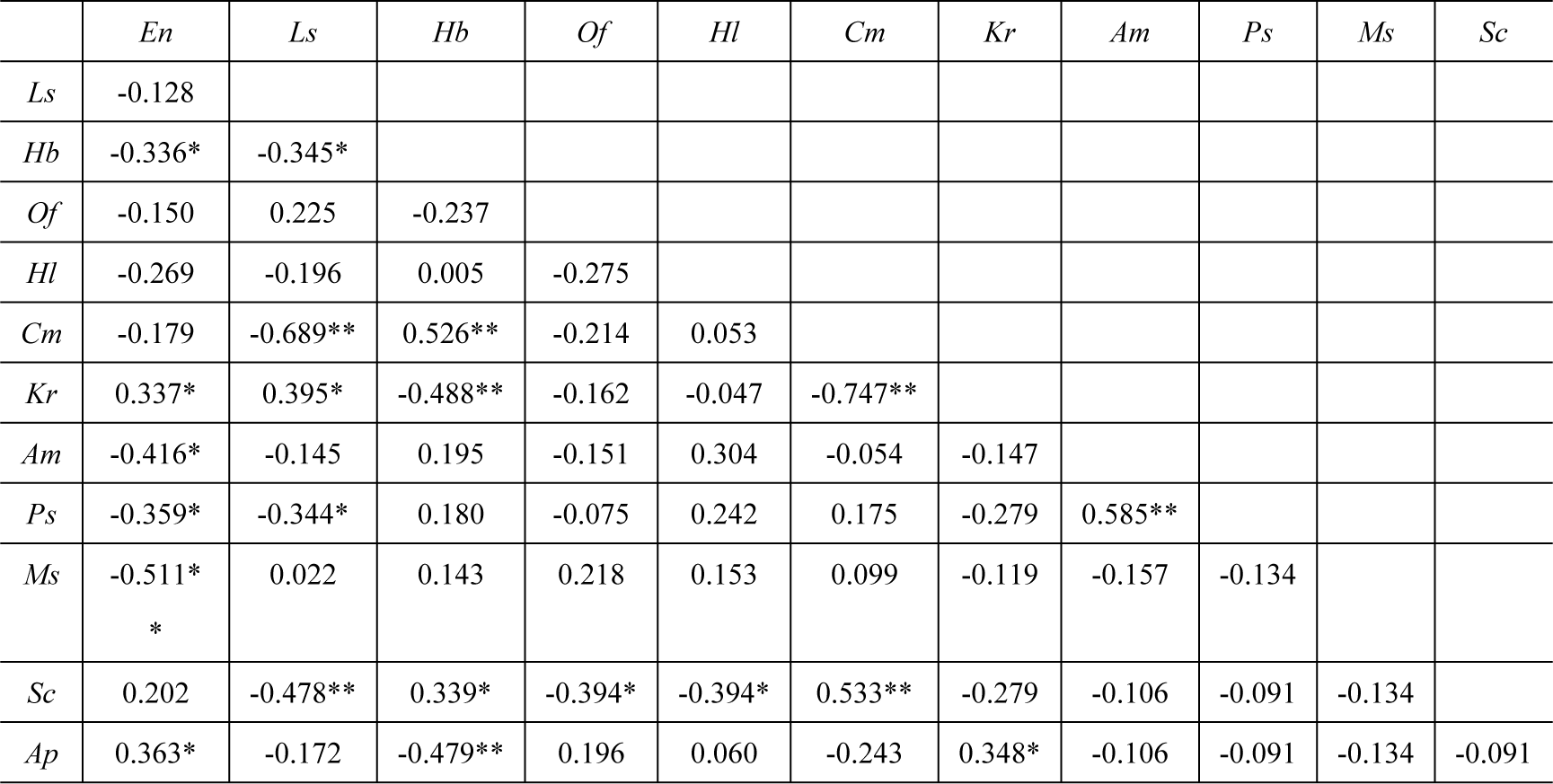
Spearman rank correlation between species-pair in wicker area.

**Table 6.** Spearman rank correlation between species-pair in control area.

|  | *En* | *Cm* | *Kr* | *Ls* | *Pb* | *Hb* | *Hl* | *Of* | *Dh* |
| --- | --- | --- | --- | --- | --- | --- | --- | --- | --- |
| *Cm* | -0.465** |  |  |  |  |  |  |  |  |
| *Kr* | 0.102 | -0.614** |  |  |  |  |  |  |  |
| *Ls* | -0.314 | -0.277 | 0.068 |  |  |  |  |  |  |
| *Pb* | -0.019 | -0.181 | 0.405* | -0.241 |  |  |  |  |  |
| *Hb* | -0.085 | 0.405* | -0.487** | -0.380* | -0.392* |  |  |  |  |
| *Hl* | 0.118 | 0.101 | -0.455** | -0.359* | -0.253 | 0.701** |  |  |  |
| *Of* | 0.278 | 0.312 | -0.436** | -0.243 | -0.172 | -0.207 | -0.134 |  |  |
| *Dh* | -0.207 | -0.326 | 0.246 | 0.596** | -0.228 | -0.275 | -0.178 | -0.121 |  |
| *Ms* | -0.376* | 0.082 | 0.027 | 0.224 | -0.172 | -0.207 | -0.134 | -0.091 | -0.121 |

Note: \*, $P < 0.05$ ; \*\*, $P < 0.01$ .

### 3.3. Niche breadth and overlap in different restoration area

Population niche breadth and niche overlap analyses could effectively assess the resources utilization and interspecific competition. The niche breadth ranged from 0.48 to 1.55 in control area, 0.38 to 1.54 in wicker area, and 0.29 to 1.55 in sandbag area. *E. nutans* had the widest niche breadth in all restoration areas. And the drought resistance plants, such as *H. bowerii*, and *C. moorcroftii* also occupied a relative wide niche breadth in all restoration areas. Furthermore, sand barriers provided a possibility for some other species’ dispersal and settle down, such as *P. sibiricum, S. collina*, and *F. ovina* (Table 2).

The average population niche overlap indices in sandbag, wicker, and control areas were 0.39, 0.32, and 0.26, respectively. Furthermore, the niche overlap indices species pair number ratio that higher than 0.50 were 33.33%, 25.76%, and 20.00% in sandbag, wicker, and control areas, respectively. The increased niche overlap indicated that the competition was stronger after the sand barriers were implemented. Moreover, there was a stronger effect of sandbag sand barriers on plant community (Figure 4).

**Figure 4.**
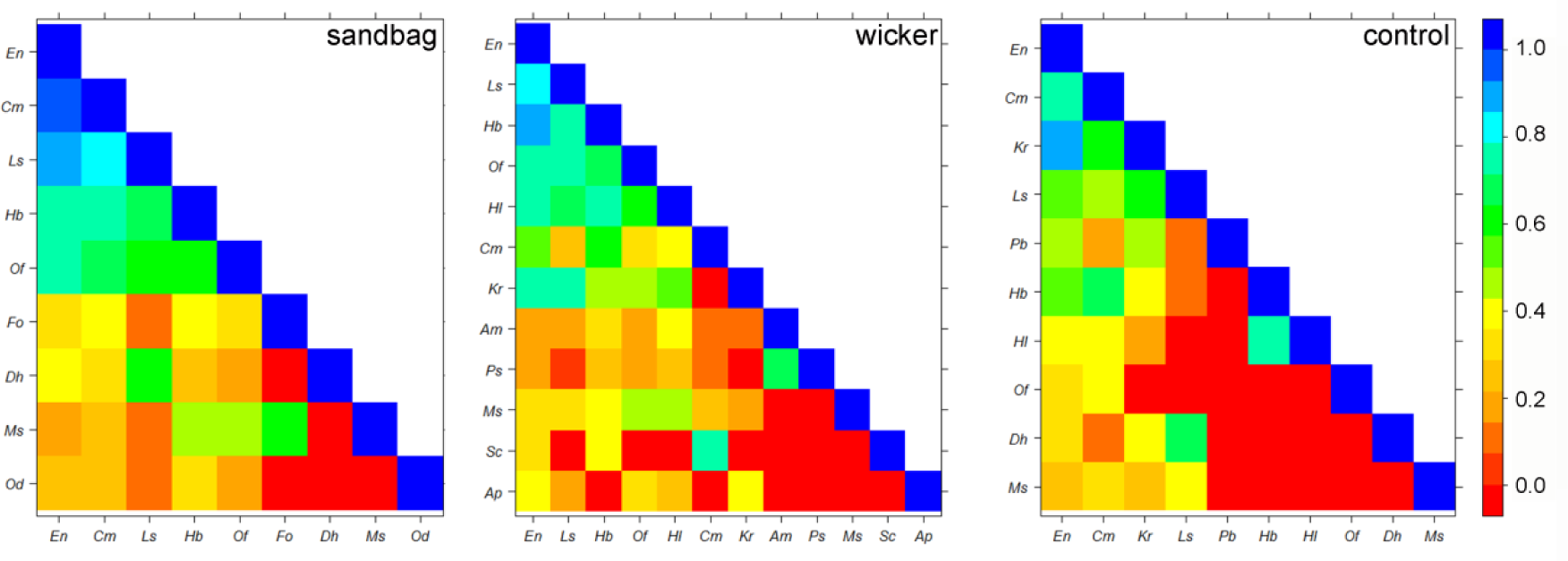
Niche overlap of all plant pairs in different restoration area.

## 4. Discussion

The sandbag and wicker sand barriers co-applied with seeding amendment were both practical approaches and were better than that only seeding during alpine sand dunes restoration. The vegetation and soil conditions were the best in the sandbag area and also were improved greatly in the wicker area compared with the control. In the control area, the soil was extremely droughty and poor, and the vegetation cover was also the lowest. The soil moisture and nutrients conditions are the preconditions that regulate the plant growth on the sand dune [36]. The soil moisture and nutrient conditions were improved greatly in the sandbag area. The sandbag is airtight whereas the wicker has a higher porosity and there was no protection in the control area. This difference leads to different near-surface wind velocities that pass by the restoration area [37, 38] (Fig 1). The strong wind would take away the soil water and destabilize the surface soil which makes it difficult for plant to sprout and grow. In addition, although the soil nutrients were was lower than the wetland where the TN was 4.9-12.0 g kg^-1^ on the Zoige Plateau [39], they were promoted greatly under wicker or sandbag amendments. The greatly increased microbial mass indicated higher microbe richness. Hence, it accelerated the litter decomposition in the soil and fed back a nutrients increasing which promoted the nitrogen content and led a lower MBC: MBN ratios [40]. The SOC: TN ratio in sandbag area also decreased to close to the level of the meadow on the Zoige Plateau (SOC: TN=11.8) [41]. Nonetheless, the SOC: TP and TN: TP ratios were far less than the ratios reported in the meadow which indicated a clear phosphorus inhibition [42]. These changes suggested that the nutrients inhibition degree was reduced, such as nitrogen inhibition was reduced, when the sand barriers were implemented. And the amendment effect of sandbag sand barrier on soil conditions was stronger than that of wicker. Moreover, soil amendments may also provide indispensable assistance during the sand dunes restoration. Except for soil nutrients regulation, bioactive fertilizer amendment may also be an effective and environmentally friendly measure to promote the microbial biomass and accelerate restoration in alpine areas [43, 44].

Plant interspecific association is important for revealing how species interact with each other and adapt with the environment, and hence have important implications for optimal restoration in degraded ecosystems [45]. Species interspecific relationships or niches play a critical role in stabilizing community [30, 46]. The tighter interspecific correlation and higher niche overlap reflected a stronger plants competitive relationship when the sand barriers were conducted [47]. The population space occupancy and correlation degree was the lowest in control while highest in sandbag area. Hence, the sandbag barrier may lead to the best plant community development in sand dune restoration [37]. The interspecific association degree was enhanced when the community was improved by wicker or sandbag barriers. However, some previous studies stated that interspecific competition/association intensity reduced gradually with the plant community development [48, 49]. The plant community that developed in alpine sand dunes area was still with limited structure and minimal resource acquisition ability, thus the independence between vascular plants was strong in such barren habitat [50].

Plant communities in sand dunes area are sensitive and vulnerable to environment changes, indicating the orientation of plant community development as well [51], allowing for revealing quantitatively community assembly mechanism or community stability [49, 52]. Plant species survived and reproduced within different restoration areas, and the population niche and interspecific relationships changed along this abiotic gradient [53]. These changes stimulated the development of the sand dune community [49, 54, 55]. Resource variations cause populations to adopt different ecological strategies to intersect with other populations [50, 56, 57]. The *E. nutans* importance value improved in the sand barrier area, especially in the sandbag area. And it occupied the wider niche to compete for the soil and light resource with the similar strategies species and coexist with the different niche requirement species. For example, *E. nutans*-*C. moorcroftii* changed from greatly negative correlation to positive correlation while the negative correlation degree of *E. nutans*-*H. bowerii* was enhanced in wicker and sandbag area. Thus, it suggested that the seeded grass regulated the relationship with other species with similar or different strategies to adapt the changed habitat and thus increase the vegetation cover. Accordingly, we also suggested that it is important to restore sand dunes by preliminarily developing a community with different ecological strategies.

## 5. Conclusions

The alpine sand dunes restoration by implementing the sand barriers and indigenous grass enhances community structure and improves the soil conditions. Using sandbag or wicker sand barriers to fix the active sand dunes would gain a better restoration effect than that only seeding. Moreover, sandbag sand barrier allowed for a better restoration of harsh soil conditions and plant community. We suggest that species interspecific relationships and niche breadth could assess the sand dune restoration efficiency well. And the soil amending measures including nutrient improvement, and microbial fertilizer addition may further accelerate sand dune restoration.

## Author Contributions

Conceptualization: Jinxing Zhou.

Data curation: Jiufu Luo.

Formal analysis: Jiufu Luo.

Funding acquisition: Jinxing Zhou, Dongzhou Deng, Li Zhang.

Investigation: Jiufu Luo, Dechao Chen, Xinwei Zhu.

Methodology: Jiufu Luo, Dongzhou Deng, Jinxing Zhou.

Project administration: Li Zhang, Dechao Chen.

Resources: Dongzhou Deng, Li Zhang, Dechao Chen, Xinwei Zhu.

Software: Jiufu Luo.

Supervision: Jinxing Zhou, Dongzhou Deng.

Validation: Jinxing Zhou, Dongzhou Deng.

Visualization: Jiufu Luo, Dechao Chen.

Writing ± original draft: Jiufu Luo.

Writing ± review & editing: Jinxing Zhou

## Acknowledgments

This study was funded by the Special Fund for Forest Scientific Research in the Public Welfare (NO. 201504401), the National Science & Technology Pillar Program (NO. 2015BAC05B01), Sichuan Province Science and Technology Support Program (NO. 2016FZ0042), and the Alpine Sand Dunes Biocrusts Composition and Characteristics on the Northwest of Sichuan (NO. 2019CZZX24).

